# A Stronger Association Between Screen Time and Externalizing Problems in Typically Developing Children than in Children with Autism Spectrum Disorder

**DOI:** 10.64898/2026.05.24.727542

**Authors:** Shogo Miyashita, Tetsu Hirosawa, Yuko Yoshimura, Chiaki Hasegawa, Sanae Tanaka, Yoshiaki Miyagishi, Nobushige Naito, Mitsuru Kikuchi

## Abstract

Excessive screen use is associated with childhood behavioral problems, but whether associations differ between typically developing (TD) children and those with autism spectrum disorder (ASD) is unclear. Our cross-sectional study included 108 children aged 5–9 years (61 TD, 47 ASD). ASD was diagnosed using standardized clinical instruments. Measures included parent-reported screen time (excluding TV/DVD), cognitive ability (K-ABC), and behavioral problems (Vineland-II). Screen time and externalizing problems were associated in the TD group (Spearman’s ρ = 0.361, p < 0.01), but not in the ASD group. In the regression model, screen time (β = 0.40, t = 2.60, p < 0.05), ASD status (β = 0.70, t = 8.30, p < 0.001), and their interaction (β = −0.34, t = −2.06, p < 0.05) significantly predicted externalizing problems. Considering the diversity within the autism spectrum, future studies with larger sample sizes should consider individual heterogeneity when examining the association between behavioral outcomes and screen time.

## Introduction

Screen exposure has become increasingly prevalent in children ^(1)^. A study involving 13,413 elementary school children (fourth to sixth grades) in Japan found that 29.9% spend more than 3 hours daily on recreational screen use ^(2)^.

The American Academy of Pediatrics (AAP) and World Health Organization (WHO) guidelines recommend screen time limits for children aged ≤ 5 years ^(3, 4)^. For children aged ≥ 6 years, the AAP guidelines recommend establishing and managing screen time rules within the family. Screen time during childhood is associated with cognitive and behavioral development, particularly those related to attention problems and externalizing behaviors. Studies involving preschoolers have reported that more than one hour of daily and early exposures (before age 2) is associated with oppositional and other problem behaviors and learning difficulties ^(5)^. Another cohort study has reported that high screen time was associated with a subsequent increase in attention problem scores and decrease in developmental test performance ^(6, 7)^. Screen time is often associated negatively with autism spectrum disorder (ASD); individuals with ASD tend to spend longer screen time than the general population ^(8)^. Children’s ASD diagnosis is associated with long hours of gameplay and problematic game use. Furthermore, high screen time is associated with a higher risk for shortened sleep duration in children with ASD ^(9, 10)^.

The prevalence of ASD diagnosis in childhood is 3.22% in Japan ^(11)^, and 2.76% in the United States ^(12)^. The mean age at which children are diagnosed with ASD is 43.18 months ^(13)^. Given the importance of early diagnosis and intervention in ASD, efforts to identify earlier diagnostic markers are accelerating ^(14, 15)^. Consequently, the mean age at diagnosis is anticipated to decline. Therefore, a more detailed understanding of the association between screen time and developmental outcomes is increasingly important for informing early support and intervention strategies.

However, few studies have examined how neurodevelopmental diagnoses relate to the association between screen time and psychological/behavioral problem scores in early school-age children (approximately 4–8 years); even fewer have directly compared the association between children who are typically developing (TD) and those with ASD. Therefore, the present study examined whether the strength of association between screen time and behavioral outcomes differs between TD and ASD groups—that is, whether there is heterogeneity by diagnosis.

## Methods

### Participants

The participants were recruited through public announcements by the Research Center for Child Mental Development, Kanazawa University. The sample included 62 TD children and 57 with ASD. ASD was diagnosed using either or both of the Autism Diagnostic Observation Schedule, Second Edition (ADOS-2) ^(16)^ and Diagnostic Interview for Social and Communication Disorders (DISCO) ^(17)^. TD children had no prior diagnosis of psychiatric or neurodevelopmental disorders and no reported problems in daily life. Children with a history of serious physical or mental illness or low birth weight were excluded. This study was approved by the Ethics Committee of the Kanazawa University Graduate School of Medical Sciences and conducted in accordance with the Declaration of Helsinki. The study period was from 22/07/2009 to 31/08/2020. Written informed consent was obtained from the parents or legal guardians of all participants.

### Measures

Screen time (excluding TV and DVD viewing) was assessed using the Japanese Sleep Questionnaire for Preschoolers ^(18)^, a parent-reported measure of children’s sleep and daily media use. The average daily screen time of the child was reported by the parents in a survey. The Kaufman Assessment Battery for Children (K-ABC) ^(19)^ and Vineland Adaptive Behavior Scales, Second Edition (Vineland-II) were administered to both TD and ASD groups ^(20)^. The K-ABC is a standardized assessment tool that uses pictures and other items to assess a child’s cognitive abilities, including processing and learning skills. The Vineland-II uses a semi-structured interview with one parent to evaluate the child’s personal and social adaptive behaviors in daily life.

### Statistical Analysis

First, t-tests were conducted to examine the group differences in demographic and psychological characteristics. Next, correlations between screen time and the Vineland-II internalizing and externalizing problems scores were examined in both groups. As the Shapiro-Wilk test indicated that screen time was not normally distributed in either the ASD group (W = 0.714, n = 47, p < 0.001) or the TD group (W = 0.584, n = 61, p < 0.001), Spearman’s rank correlation coefficient (ρ) was used for the correlation analyses.

To reduce the likelihood of Type I errors due to multiple testing, the Bonferroni correction was applied to control the family-wise error rate. Considering two groups (ASD and TD) and two correlation tests (internalizing and externalizing problems raw scores), the significance threshold was set at α = 0.05/4 = 0.0125.

Spearman’s rank correlation analysis revealed a significant association only between screen time and the Vineland-II externalizing problems score in the TD group; therefore, the subsequent analyses focused on this outcome. A multiple linear regression analysis was performed to examine whether the association between screen time and Vineland-II externalizing problems scores differed between the groups. In this model, the Vineland-II externalizing problems score was entered as the dependent variable, whereas screen time, ASD status (TD = 0, ASD = 1), and their interaction term were entered as independent variables.

## Results

### Participant Characteristics

One TD child and one with ASD were excluded due to missing screen time data. To standardize the intellectual ability level across both groups, children with a K-ABC mental processing score less than 70 were excluded (0 in TD, 9 in ASD; Figure 1). Participant characteristics are summarized in Table 1. Primary Analyses Screen time was significantly positively correlated with Vineland-II externalizing problems scores in the TD group (n = 61; Spearman’s ρ = 0.361, p < 0.01). No correlation was observed in the ASD group (ρ = 0.024, p = 0.873). Furthermore, neither group showed a significant correlation between screen time and internalizing problem scores (TD: ρ = 0.149, p = 0.252; ASD: ρ = 0.104, p = 0.485). Figure 2 shows the relationship between screen time and each score in the TD group.

**Table 1.** Demographic characteristics of the study participants (mean± standard deviation or%)

|  | TD | ASD | TD vs. ASD |
| --- | --- | --- | --- |
| Number of participants | 61 | 47 | - |
| Sex (male / female) | 40 / 21 | 32 / 15 | - |
| Chronological age in months [range] (SD) | 70.72 [60-92]<br>(9.69) | 74.51 [60-111]<br>(12.36) | t = -1.79 |
| K-ABC mental processing score (SD) | 108.43 (14.38) | 96 (17.41) | t = 4.06** |
| Screen time (SD) | 15.25 (29.83) | 40.53 (61.2) | t = -2.6* |

Note. Group differences were examined using t-tests. TD = typically developing; ASD = autism
Battery for Children. \*p<0.05, \*\*p<0.001

**Figure 1.**
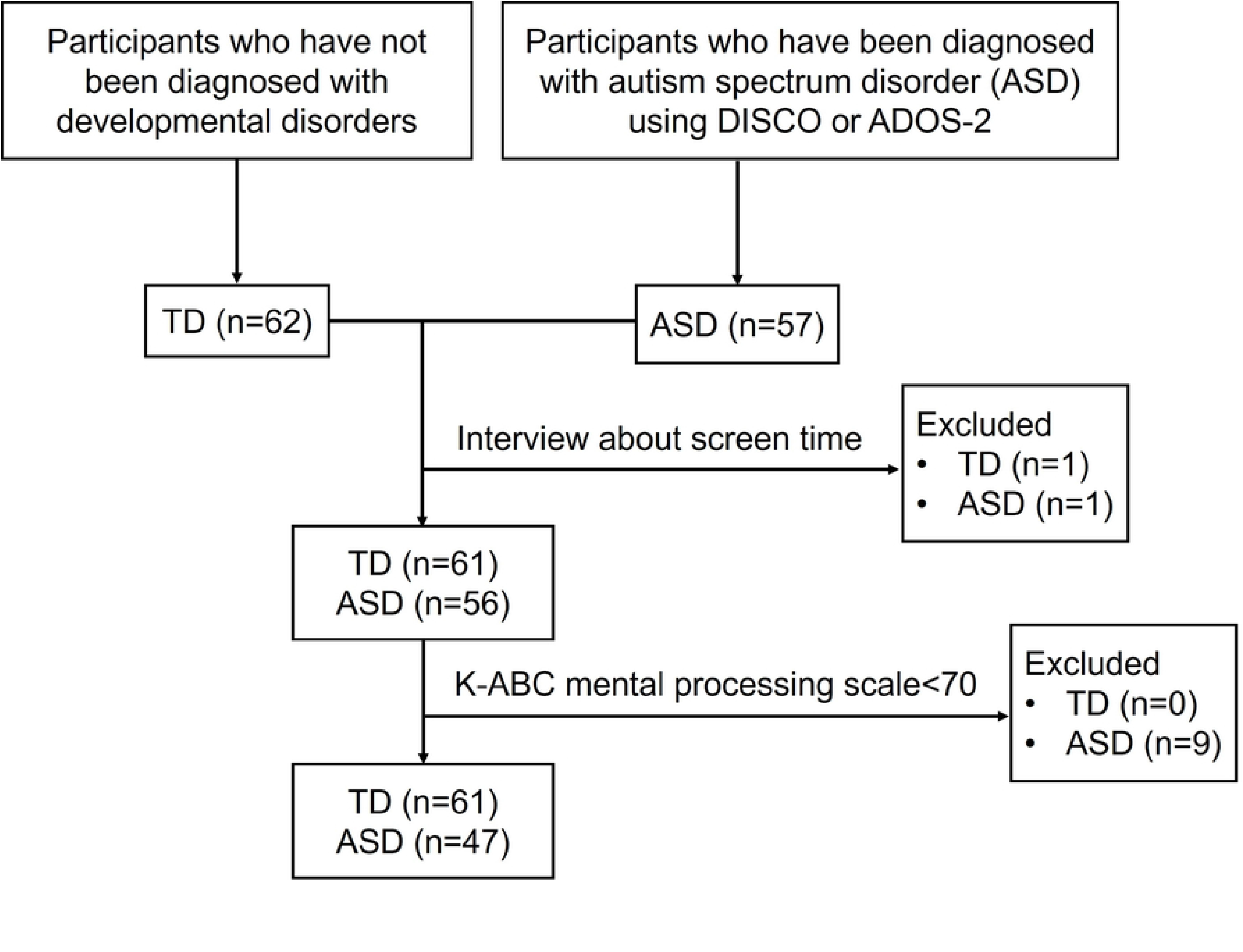
Flow chart of participant recruitment and inclusion. Note. This figure shows the recruitment process and the number of participants included in this study. TD = typically developing; ASD = autism spectrum disorder; ADOS-2 = Autism Diagnostic Observation Schedule, Second Edition; DISCO = Diagnostic Interview for Social and Communication Disorders; K-ABC = Kaufman Assessment Battery for Children. The figure also shows exclusions due to missing screen time data and K-ABC mental processing scores < 70.

**Figure 2.**
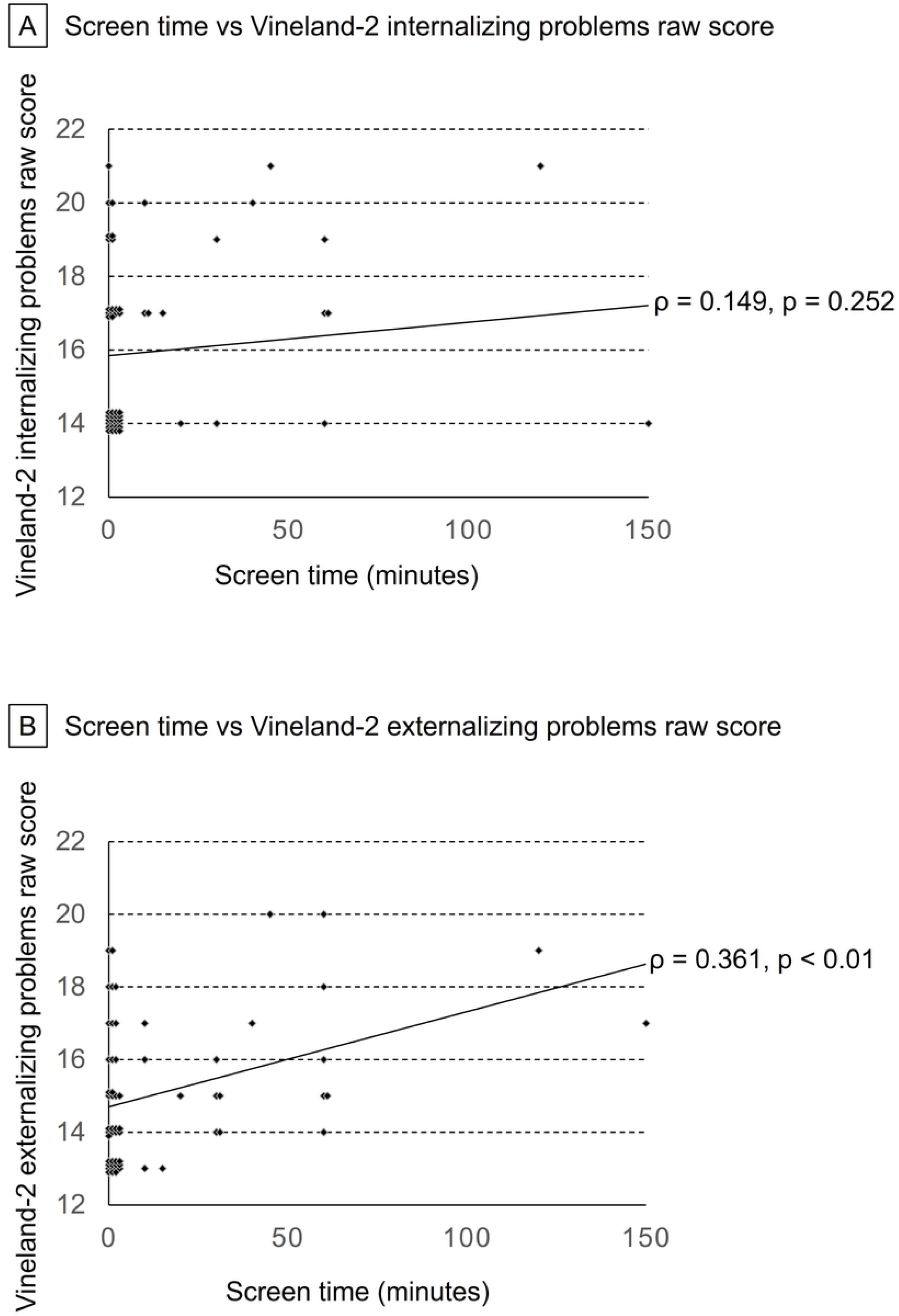
Scatter plot showing the relationship between screen time and problem behavior scores in typically developing children. Note. The two graphs illustrate the relationship between screen time and the Vineland-II internalizing and externalizing problems scores in the typically developing group. Screen time was significantly positively correlated with the Vineland-II externalizing problems score in the TD group (n = 61; Spearman’s ρ = 0.361, p < 0.01), but not with internalizing problems score.

### Secondary Analyses

The initial results suggested that the relationship between screen time and externalizing behavioral problems differed between the TD and ASD groups. To test this, a multiple linear regression analysis (n = 108) was performed. In the multiple linear regression analysis, screen time (β = 0.40, t = 2.60, p < 0.05), ASD status (β = 0.70, t = 8.30, p < 0.001), and their interaction term (β = −0.34, t = −2.06, p < 0.05) significantly predicted the Vineland-II externalizing problems score (Table 2). The model was significant (F value = 29.58, p < 0.001), with a robust fit, yielding a coefficient of determination of R^2^ = 0.46 (adjusted R2 = 0.45). The significant interaction term in the model indicated that the association between screen time and the externalizing problems score differed between the groups. Collectively, the primary analyses demonstrated a positive correlation between screen time and Vineland-II externalizing problems in the TD group. The subsequent regression model, which incorporated the interaction term, confirmed that the association between screen time and the Vineland-II externalizing problems score differed significantly between the TD and ASD groups (Figure 3).

**Table 2.** Results of multiple regression analysis predicting Vineland-II externalizing problems scores.

| | $\beta$ |
| --- | --- |
| Screen time (minutes) | 0.400* |
| ASD or TD (ASD = 1, TD = 0) | 0.699** |
| Interaction term | -0.342* |
| F-value | 29.582** |
| R-value | 0.679 |
| Adjusted R<sup>2</sup> | 0.445 |

Note. The model included screen time, ASD status (ASD = 1, TD = 0), and the interaction term
(screen time × ASD status) as predictors. TD = typically developing; ASD = autism spectrum
disorder; ADOS-2 = Autism Diagnostic Observation Schedule, Second Edition; DISCO = Diagnostic
Interview for Social and Communication Disorders; K-ABC = Kaufman Assessment Battery for
Children; Vineland-II = Vineland Adaptive Behavior Scales, Second Edition. \* $p < 0.05$ \*\* $p < 0.001$

**Figure 3.**
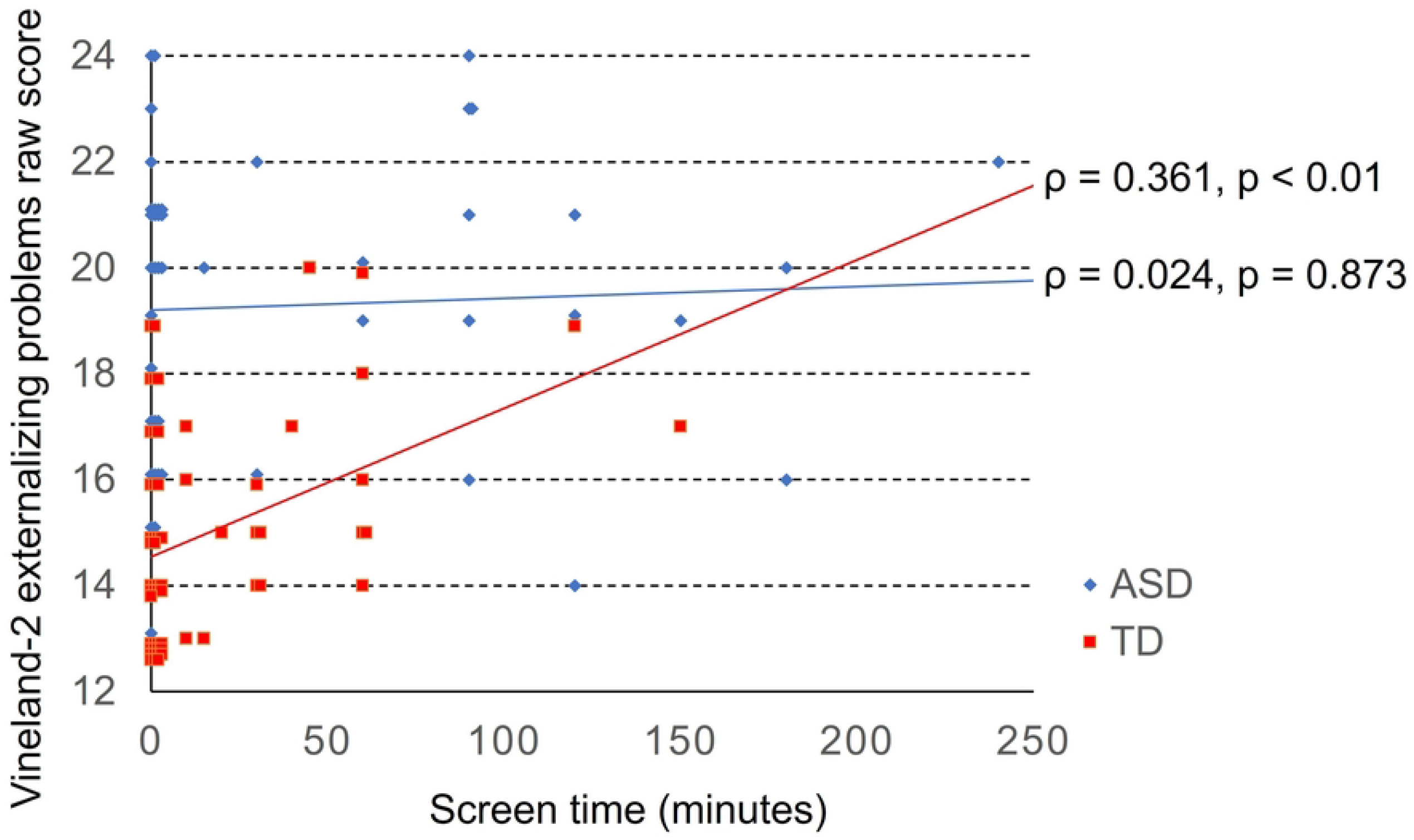
Scatter plot of screen time and Vineland-II externalizing problems raw score in both groups. Note. TD = typically developing; ASD = autism spectrum disorder. The TD and ASD groups are plotted separately to visualize the differences in association between diagnostic groups. In the multiple linear regression analysis, screen time (β = 0.40, t = 2.60, p < 0.05), ASD status (β = 0.70, t = 8.30, p < 0.001), and their interaction term (β = −0.34, t = −2.06, p < 0.05) significantly predicted the Vineland-II externalizing problems score.

## Discussion

This study observed that children with ASD have longer screen time than TD children, consistent with prior reports documenting greater screen use in children with ASD ^(21, 22)^. In the primary analyses, Vineland-II internalizing problems scores were not significantly associated with screen time in either group. This pattern accords with follow up studies suggesting that the associations between screen time and internalizing behaviors in preschool years may change around school entry ^(23)^. Large scale studies have reported small but statistically significant associations between higher screen time and internalizing problems, with smaller effect sizes than in those for externalizing behaviors ^(24)^. Accordingly, the present sample may have been underpowered to detect such associations.

By contrast, the Vineland-II externalizing problems score was positively associated with screen time in the TD group, but not in the ASD group. The literature on screen use in ASD reports no significant associations between screen time frequency or duration—particularly game play—and aggression or irritability ^(25)^, which is consistent with our ASD group findings. Some intervention studies have reported ASD symptom improvement and parenting stress reduction following efforts to reduce screen exposure ^(26, 27)^; however, most of these studies used small samples and heterogeneous designs, limiting causal inferences and conclusions.

In the multiple linear regression model, the interaction between ASD status and screen time significantly predicted Vineland-II externalizing problems score, indicating that the strength of the association differed by diagnostic group. In this study, TD children exhibited a stronger positive association between screen time and externalizing problem scores than those with ASD. Observational works in similar-aged cohorts have documented that reduced screen time periods coincided with lower aggression and delinquency, including fewer verbal aggression incidents at school ^(28, 29)^. Nevertheless, such converging observations cannot determine directionality; children with higher externalizing tendencies may engage in greater screen use, greater screen use may co-occur with higher externalizing scores or shared third variables may contribute to both.

Potential mechanisms remain speculative. Neuroimaging studies have reported reduced volumes of the dorsolateral prefrontal cortex, orbitofrontal cortex, and insular cortex and lower striatal dopamine receptor levels in individuals with internet gaming disorder compared with healthy controls ^(30)^. Greater screen use is associated with lower structural integrity of the uncinate fasciculus and inferior longitudinal fasciculus in preschoolers ^(31)^. While these findings support the differences in neural systems relevant to behavioral control, they are correlational and do not establish causation.

This study has several limitations. Screen time and Vineland-II behavioral indicators were reported by parents and are susceptible to recall and social desirability biases. Children with ASD typically present with higher baseline problem behavior scores, which could attenuate detectable associations with screen time in this group. In addition, this study did not distinguish passive from active screen time, despite evidence that passive exposure is associated with poorer verbal processing and self-regulation, whereas active screen time has been associated with better working memory and receptive language skills ^(32)^. Differentiating these modalities may allow more precise estimation of the associations with behavioral scores.

The present study focused on school-age children and did not evaluate interventions. However, the observed pattern indicates that screen time and externalizing problems scores were positively associated in TD children but not in those with ASD. Moreover, higher screen time and problem behavior scores were not significantly associated in children with ASD. However, given the cross-sectional design and limited sample size, the absence of an association in the ASD group should not be interpreted as evidence of no impact of screen time on behavioral outcomes in ASD, and no causal conclusions can be drawn regarding the direction of the associations observed in either group. Thus, the present cross-sectional findings do not establish that limiting screen time would reduce behavioral problems in this sample. Future large-scale, prospective studies that account for heterogeneity within the autism spectrum and distinguish passive from active media are needed to clarify these associations and their potential relevance for early support strategies.

## Notes

### Competing Interest Statement

The authors have declared no competing interest.

## References

1. Madigan S, Eirich R, Pador P, McArthur BA, Neville RD. Assessment of Changes in Child and Adolescent Screen Time During the COVID-19 Pandemic: A Systematic Review and Meta-analysis. JAMA Pediatr. 2022;176(12):1188–98.

2. Yamada M, Sekine M, Tatsuse T. Association between excessive screen time and school-level proportion of no family rules among elementary school children in Japan: a multilevel analysis. Environ Health Prev Med. 2024; 29:16.

3. Council On C, Media. Media Use in School-Aged Children and Adolescents. Pediatrics. 2016;138(5).

4. WHO Guidelines Approved by the Guidelines Review Committee. Guidelines on Physical Activity, Sedentary Behaviour and Sleep for Children under 5 Years of Age, 2019’. Geneva: World Health Organization.; 2019.

5. Xiang H, Lin L, Chen W, Li C, Liu X, Li J, et al. Associations of excessive screen time and early screen exposure with health-related quality of life and behavioral problems among children attending preschools. BMC Public Health. 2022;22(1):2440.

6. Swing EL, Gentile DA, Anderson CA, Walsh DA. Television and video game exposure and the development of attention problems. Pediatrics. 2010;126(2):214–21.

7. Madigan S, Browne D, Racine N, Mori C, Tough S. Association Between Screen Time and Children’s Performance on a Developmental Screening Test. JAMA Pediatr. 2019;173(3):244–50.

8. Ophir Y, Rosenberg H, Tikochinski R, Dalyot S, Lipshits-Braziler Y. Screen Time and Autism Spectrum Disorder: A Systematic Review and Meta-Analysis. JAMA Netw Open. 2023;6(12):e2346775.

9. Mazurek MO, Engelhardt CR. Video game use in boys with autism spectrum disorder, ADHD, or typical development. Pediatrics. 2013;132(2):260–6.

10. Engelhardt CR, Mazurek MO, Sohl K. Media use and sleep among boys with autism spectrum disorder, ADHD, or typical development. Pediatrics. 2013;132(6):1081–9.

11. Saito M, Hirota T, Sakamoto Y, Adachi M, Takahashi M, Osato-Kaneda A, et al. Prevalence and cumulative incidence of autism spectrum disorders and the patterns of co-occurring neurodevelopmental disorders in a total population sample of 5-year-old children. Mol Autism. 2020;11(1):35.

12. Maenner MJ, Warren Z, Williams AR, Amoakohene E, Bakian AV, Bilder DA, et al. Prevalence and Characteristics of Autism Spectrum Disorder Among Children Aged 8 Years - Autism and Developmental Disabilities Monitoring Network, 11 Sites, United States, 2020. MMWR Surveill Summ. 2023;72(2):1–14.

13. van ‘t Hof M, Tisseur C, van Berckelear-Onnes I, van Nieuwenhuyzen A, Daniels AM, Deen M, et al. Age at autism spectrum disorder diagnosis: A systematic review and meta-analysis from 2012 to 2019. Autism. 2021;25(4):862–73.

14. Wallace KS, Rogers SJ. Intervening in infancy: implications for autism spectrum disorders. J Child Psychol Psychiatry. 2010;51(12):1300–20.

15. Zwaigenbaum L, Bauman ML, Stone WL, Yirmiya N, Estes A, Hansen RL, et al. Early Identification of Autism Spectrum Disorder: Recommendations for Practice and Research. Pediatrics. 2015;136 Suppl 1(Suppl 1): S10–40.

16. Lord C, Western Psychological S. ADOS-2: autism diagnostic observation schedule: manual. 2nd ed. Los Angeles, Calif: Western Psychological Services; 2012.

17. Wing L, Leekam SR, Libby SJ, Gould J, Larcombe M. The Diagnostic Interview for Social and Communication Disorders: background, inter-rater reliability and clinical use. J Child Psychol Psychiatry. 2002;43(3):307–25.

18. Shimizu S, Kato-Nishimura K, Mohri I, Kagitani-Shimono K, Tachibana M, Ohno Y, et al. Psychometric properties and population-based score distributions of the Japanese Sleep Questionnaire for Preschoolers. Sleep Med. 2014;15(4):451–8.

19. Kaufman AS, Kaufman NL. K-ABC: Kaufman Assessment Battery for Children: Interpretive Manual: American Guidance Service; 1983.

20. Sara SS, Domenic C, David AB. Vineland Adaptive Behavior Scales, Second Edition. American Psychological Association (APA); 2005.

21. Hill MM, Gangi DN, Miller M. Toddler Screen Time: Longitudinal Associations with Autism and ADHD Symptoms and Developmental Outcomes. Child Psychiatry Hum Dev. 2024.

22. Takahashi N, Tsuchiya KJ, Okumura A, Harada T, Iwabuchi T, Rahman MS, et al. The association between screen time and genetic risks for neurodevelopmental disorders in children. Psychiatry Res. 2023; 327: 115395.

23. Neville RD, McArthur BA, Eirich R, Lakes KD, Madigan S. Bidirectional associations between screen time and children’s externalizing and internalizing behaviors. J Child Psychol Psychiatry. 2021;62(12):1475–84.

24. Eirich R, McArthur BA, Anhorn C, McGuinness C, Christakis DA, Madigan S. Association of Screen Time With Internalizing and Externalizing Behavior Problems in Children 12 Years or Younger: A Systematic Review and Meta-analysis. JAMA Psychiatry. 2022;79(5): 393–405.

25. Alkhayat LS, Ibrahim M. Assessing the effect of playing games on the behavior of ASD and TD children. Advances in Autism. 2020;6(4):315–34.

26. Heffler KF, Frome LR, Garvin B, Bungert LM, Bennett DS. Screen time reduction and focus on social engagement in autism spectrum disorder: A pilot study. Pediatr Int. 2022;64(1): e15343.

27. Sadeghi S, Pouretemad H, Khosrowabadi R, Fathabadi J, Nikbakht S. Behavioral and electrophysiological evidence for parent training in young children with autism symptoms and excessive screen-time. Asian J Psychiatr. 2019; 45: 7–12.

28. Yilmaz G, Demirli Caylan N, Karacan CD. An intervention to preschool children for reducing screen time: a randomized controlled trial. Child Care Health Dev. 2015;41(3):443–9.

29. Robinson TN, Wilde ML, Navracruz LC, Haydel KF, Varady A. Effects of reducing children’s television and video game use on aggressive behavior: a randomized controlled trial. Arch Pediatr Adolesc Med. 2001;155(1):17–23.

30. Zhu Y, Zhang H, Tian M. Molecular and functional imaging of internet addiction. Biomed Res Int. 2015; 2015:378675.

31. Hutton JS, Dudley J, Horowitz-Kraus T, DeWitt T, Holland SK. Associations Between Screen-Based Media Use and Brain White Matter Integrity in Preschool-Aged Children. JAMA Pediatr. 2020;174(1): e193869.

32. Bal M, Kara Aydemir AG, Tepetas Cengiz GS, Altindag A. Examining the relationship between language development, executive function, and screen time: A systematic review. PLoS One. 2024;19(12):e0314540.

